# Simultaneous Optimization of Excitation Wavelength and Emission Window for Semiconducting Polymer Nanoparticles to Improve Imaging Quality

**DOI:** 10.1101/2021.12.26.474182

**Authors:** Dingwei Xue, Hongli Zhou, Zeyi Lu, Yuhuang Zhang, Mengyuan Li, Zhousu Xu, Abudureheman Zebiluba, Zhe Feng, Lin Li, Jie Liu, Jun Qian, Gonghui Li

**Affiliations:** Department of Urology, Sir Run-Run Shaw Hospital, School of Medicine, Zhejiang University, Hangzhou, 310016, China; Key Laboratory of Flexible Electronics (KLOFE) Institute of Advanced Materials (IAM), Nanjing Tech University, Nanjing 211800, China; State Key Laboratory of Modern Optical Instrumentations, Centre for Optical and Electromagnetic Research, College of Optical Science and Engineering, International Research Center for Advanced Photonics, Zhejiang University, Hangzhou, 310058, China; Institute of Intelligent Optoelectronic Technology, Zhejiang University of Technology, Hangzhou, 310014, China

**Keywords:** semiconducting polymer nanoparticle, fluorescence imaging, near-infrared IIb window, angiography, tumor imaging

## Abstract

Optimized excitation wavelength and emission window are essential for fluorescence imaging with high quality. Semiconducting polymer nanoparticles (SPNs) as fluorescent contrast agents have been extensively studied, but their imaging abilities in the second near-infrared IIb window (NIR-IIb, 1500–1700 nm) with long excitation wavelength have not been reported yet. Herein, as a proof-of-concept, we demonstrate for the first time that an SPN named L1057 nanoparticles (NPs) exhibit intense NIR-IIb signal due to its ultra-high brightness and broad emission spectrum. After screening 915 nm as an optimal excitation wavelength, we applied L1057 NPs to visualize the whole-body vessels, cerebral vessels, gastrointestinal tract, and tumor progression in different stages, achieving superior spatial resolution and signal to background ratio in the NIR-IIb window with respect to NIR-II window (1000–1700 nm). This study reveals that simultaneous optimization of excitation wavelength and emission window is an efficient strategy to enhance imaging quality and that L1057 NPs can serve as a promising NIR-IIb contrast agent for high-resolution and deep-tissue imaging.

## 1. Introduction

Fluorescence imaging in the second near-infrared window (NIR-II, 1000□1700 nm) shows deeper tissue penetration depth and higher signal-to-background ratio (SBR) than that in the conventional near-infrared window (NIR-I, 700–900 nm), and has emerged as a promising imaging method for visualization of deep tissues.^[1]^ The first-in-human liver-tumor surgery guided by NIR-II fluorescence imaging further highlighted its great potential in preclinical and clinical practices. Optimization of excitation wavelength and emission regions is of significance to improve imaging quality. On the one hand, the excitation light traveling through biological tissues is attenuated by the absorption and scattering of tissues,^[2]^ bathochromic shift of excitation wavelength and bypassing the overtone peaks of water can improve the imaging penetration and resolution. However, most of the previously reported NIR-II fluorophores were excited within NIR-I window (e.g., 808 nm). On the other hand, the fluorescence imaging in the NIR-IIb subwindow (1500–1700 nm) has been reported to show notably enhanced spatial resolution and SBR compared to NIR-II and NIR-IIa (1300–1400 nm) imaging owing to near-zero autofluorescence and remarkably reduced photon scattering.^[3]^ The currently available NIR-IIb fluorophores are mainly limited to inorganic materials, such as single-walled carbon nanotubes (SWNTs), ^[4]^ rare-earth-doped nanoparticles (RENPs),^[5]^ and quantum dots (QDs).^[6]^ On account of unknown long-term cytotoxicity concerns of these inorganic nanomaterials, it is urgent to develop organic fluorophores to facilitate clinical translation. Until recently, organic fluorophores based on aggregation-induced emission (AIE) molecules,^[7]^ *J*-aggregates of cyanine dye^[8]^ and indocyanine green^[1c]^ exhibited NIR-IIb emission tail and have been utilized for *in vivo* NIR-IIb bioimaging. However, they generally required high dose, strong laser power density or long exposure time during imaging due to the low effective NIR-IIb brightness. Thus, the development of bright NIR-IIb fluorophores is still challenging.

Semiconducting polymer nanoparticles (SPNs) represent another important fluorescent nanomaterial used in the bioimaging field due to their high extinction coefficient, large Stokes shift, good photostability as well as excellent biocompatibility.^[9]^ Some NIR-I excitable SPNs have been reported to have NIR-II fluorescence, but NIR-IIb fluorescent SPNs have not been reported yet. Considering that SPNs generally have broad emission spectra, we speculated that highly bright SPNs may have NIR-IIb emission tail.

Recently, we reported nanoparticles, L1057 NPs, based on a semiconducting polymer PTQ. Taking advantage of rational molecular design and long conjugation length, L1057 NPs not only had longer excitation wavelength (e.g., 980 nm) but also exhibited much higher effective NIR-II brightness than many reported organic NIR-II fluorophores.^[10]^ Thus, as a proof-of-concept, we herein demonstrated the capability of L1057 NPs as NIR-IIb fluorophore for *in vivo* deep-tissue imaging. After screening 915 nm as the optimal excitation wavelength, we applied L1057 NPs as NIR-IIb contrast agent to realize *in vivo* dynamic vascular structure imaging, cerebrovascular functional imaging, gastrointestinal imaging, and tumor progression imaging in small animals, exhibiting higher spatial resolution and better SBR for bioimaging in the NIR-IIb window compared with those in the conventional NIR-II and NIR-IIa windows. To the best of our knowledge, this is the first report using SPNs as NIR-IIb contrast agent.

**Table of Contents.**
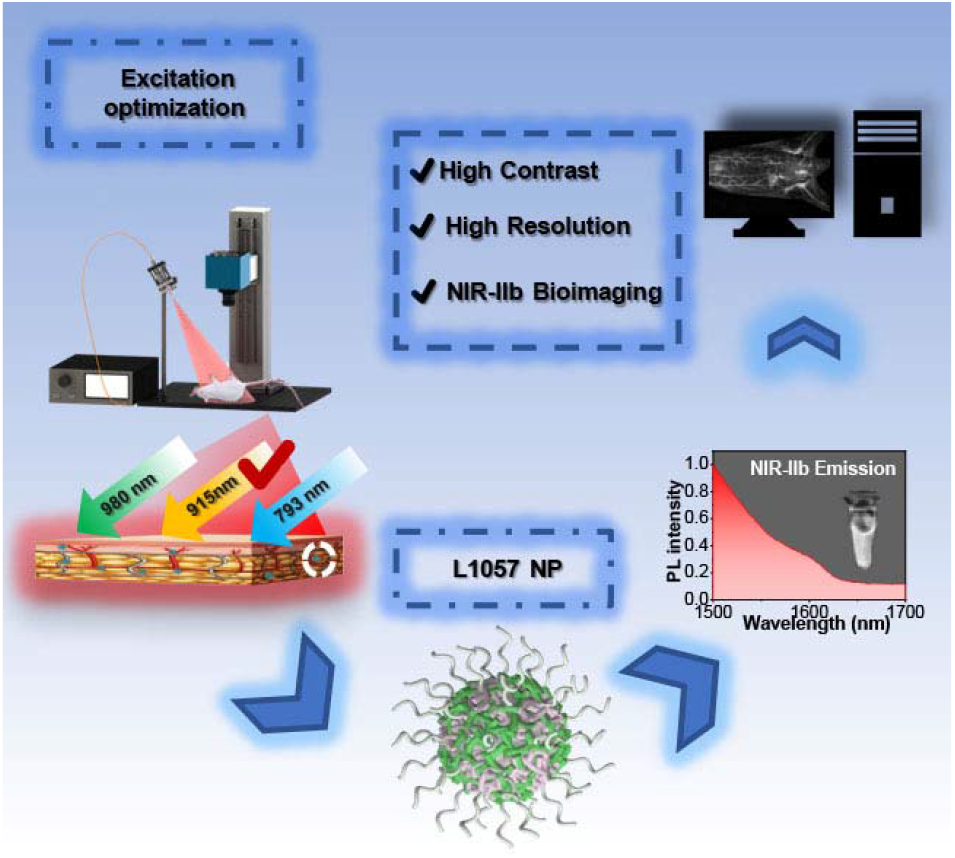
Excitation wavelength screening and high-quality NIR-IIb bioimaging of semiconducting polymer nanoparticle L1057 NP

## 2. Results and Discussion

### 2.1. Characterization of L1057 NPs

Figure 1a shows the chemical structure of PTQ and the schematic structure of L1057 NPs. PTQ was synthesized according to our previous report,^[10]^ through Pd-catalyzed Stille polymerization from 4,9-dibromo-6,7-bis(4-(hexyloxy)phenyl)-[1,2,5]thiadiaz-olo[3,4-*g*]quinoxaline and 6,6,12,12-tetrakis(4-hexylphenyl)-*s*-indacenodithieno[3,2-*b*]thiophene-bis(trimethylst annane).

**Figure 1.**
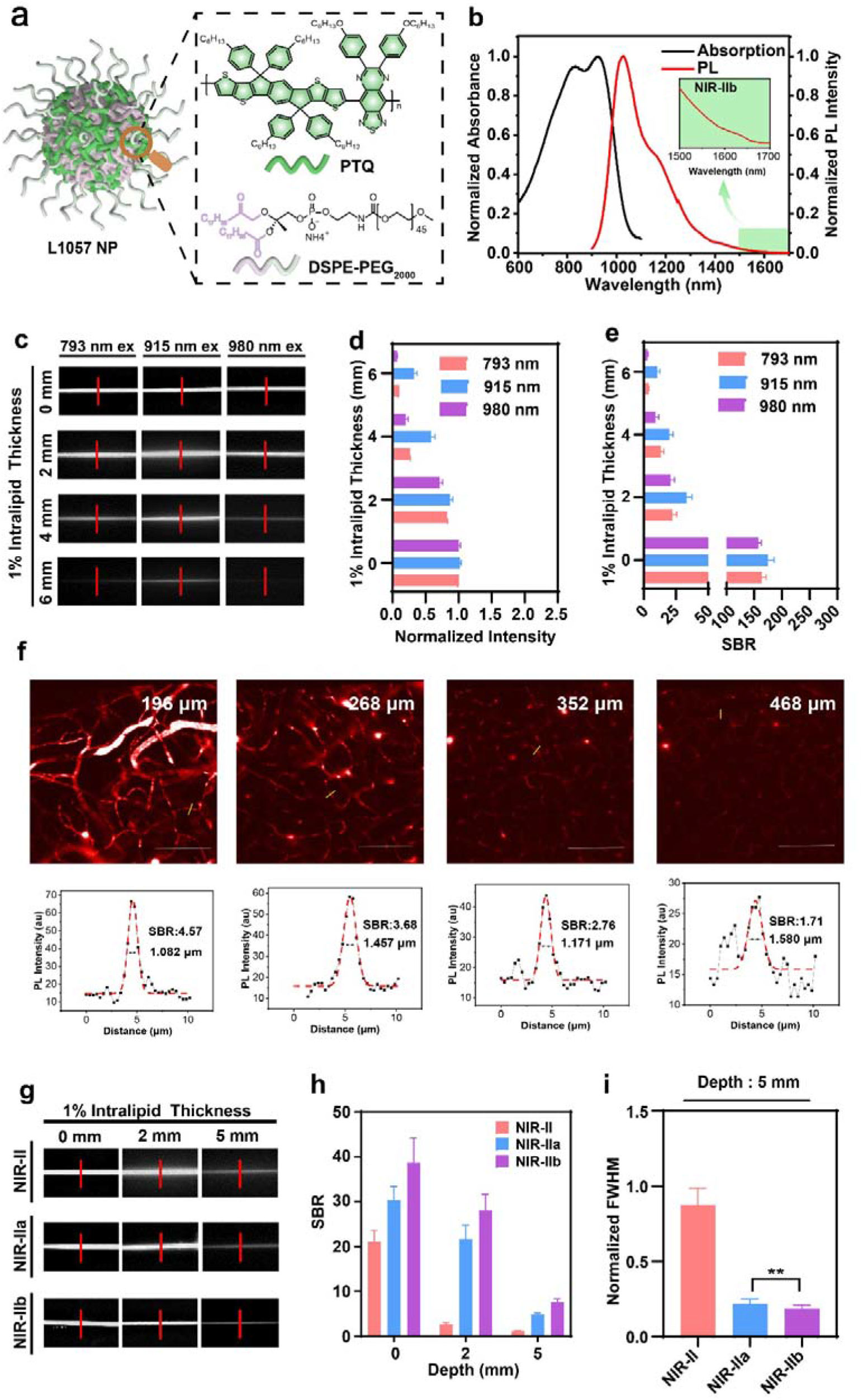
*In vitro* characterization of L1057 NPs. (a) Schematic of L1057 NPs and the chemical structures of PTQ and DSPE-PEG_2000_. (b) Normalized absorption and emission spectra of L1057 NPs in deuteroxide. (c) The fluorescence images of L1057 NPs (1 mg mL^-1^)-loaded capillary glass tube embedded in Intralipid® solution, with excitation at 793, 915 and 980 nm (5 mW cm^-2^), respectively, and signal collection using a 1200 nm LP filter. (d) Quantitative analysis of fluorescence intensity attenuation and (e) SBRs as a function of Intralipid® thickness upon different excitation wavelength. Data are the mean ± SD, n = 3. (f) NIR-II fluorescence confocal microscopic in vivo imaging of cerebral blood vessels of the mouse with high lateral resolution and SBR at 4 typical depths (196 μm, 268 μm, 352 μm and 468 μm). Yellow lines: locations of cross-sections depicted in upper row. The cross-sectional fluorescence intensity profiles (black) and the related Gaussian fits (red) along the capillary vessels, taken from locations (yellow lines) in the lower row. Excitation wavelength: 915 nm. Laser power: ∼60 mW before the objective. PMT voltage: ∼650 V. Pinhole diameter: 200 μm. All scale bars: 50 μm. (g) The fluorescence images of L1057 NPs (1 mg mL^-1^)-loaded capillary glass tube embedded in Intralipid® solution, with excitation at 915 nm and signal collection in different detection subwindows. (h) Quantitative analysis of SBR and (i) FWHM as a function of Intralipid® thickness in different detection subwindows. Data are the mean ± SD, n = 3.

As is illustrated in our previous study, we used a weaker electron acceptor for PTQ but the polymer still exhibited red-shifted absorption profiles owing to the longer conjugation length of SPNs relative to small molecules.^[10]^ PTQ was then encapsulated into nanoparticles to afford L1057 NPs using DSPE-PEG_2000_ as matrix to increase its biocompatibility. L1057 NPs exhibited an absorption plateau in the region of 910□940 nm with a high mass extinction coefficient (∼23 L g^-1^cm^-1)^, and a strong emission peaked at 1057 nm with a quantum yield of 1.25% in the NIR-II region (Figure 1b). As shown in the inset of Figure 1b, L1057 NPs had a long emission tail extending to 1700 nm, and a bright NIR-IIb image can be obtained upon excitation at 915 nm. To highlight the NIR-IIb properties of L1057 NPs, we compared L1057 NPs with previously reported NIR-IIb fluorophores, TT3-*o*CB NPs and indocyanine green (ICG), using their respective appropriate excitation wavelength with the same laser power. The results showed that L1057 NPs exhibited a stronger NIR-IIb fluorescence intensity than TT3-*o*CB NPs and ICG (see Figure S1). These results indicated that L1057 NP was an excellent NIR-IIb fluorophore candidate for NIR-IIb bioimaging.

To determine an optimal excitation wavelength for better imaging quality, we first performed phantom assay in 1% Intralipid® using 793, 915 and 980 nm, respectively, as excitation wavelength with the same power density (5 mW cm^-2^), and compared their normalized intensity and SBR. We chose these three excitation wavelengths based on the consideration: i) 793 and 980 nm are two commonly used excitation wavelengths for NIR-II imaging.^[1b, 1d, 2a, 3c, 5b, 7a, 10-11]^ and ii) L1057 NPs have a high mass extinction coefficient at 915 nm, which is also far away from the overtone peak of water. The fluorescence signal was collected using a 1200 nm long-pass (LP) filter. The fluorescence intensity was normalized and the SBR was analyzed at different depths and different detection subwindows. With the increase of penetration depth, the fluorescence intensity and SBR gradually decreased but the capillary excited at 915 nm showed the highest fluorescence intensity and SBR. The capillary can still be clearly distinguished even at a depth of 6 mm (Figure 1c-e). The higher fluorescence intensity and SBR obtained by excitation at 915 nm relative to the other two wavelengths could be explained by the fact that the water absorption intensity at 915 nm is 3.6-fold lower than that at 980 nm (see Figure S2) although light scattering by tissue is inversed with wavelength. On the other hand, considering that L1057 NPs have higher extinction coefficient at 915 nm than at 793 and 980 nm, 915 nm should be an appropriate excitation wavelength for L1057 NPs to obtain high-quality fluorescence imaging. Therefore, *in vivo* whole-body imaging, which was also performed with different excitation wavelength (793, 915 and 980 nm) and imaging windows (NIR-II and NIR-IIb), consistently showed that imaging in the NIR-IIb window with excitation at 915 nm revealed remarkably clearer and more detailed vessels in deep tissues compared with those with other experimental conditions (see Figure S3). We also utilized 915 nm as the optimized excitation laser to achieve excellent NIR-II fluorescence confocal microscopy of cerebral blood vessels of the mouse with fine optical sectioning and high SBR. Figure 1f exhibited representative confocal images taken at depth of 196 μm, 268 μm, 352 μm and 300 μm (upper row), respectively, and profiles and Gaussian fits of fluorescence intensity profiles along the yellow lines are shown in lower row. Besides, the penetration depth was as deep as 600 μm shown in Figure S4.All the subsequent imaging experiments in this study used 915 nm as excitation wavelength unless otherwise specified.

As L1057 NPs had an emission tail extending to the NIR-IIb region, we subsequently tried to investigate their feasibility as NIR-IIb fluorescence agent for imaging using intralipid® phantom assay. The fluorescence signals decreased with the increasing thickness of 1% Intralipid® solution (Figure 1g). The SBRs for L1057 NPs in the NIR-II and NIR-IIa window were significantly lower than that in the NIR-IIb window at different 1% Intralipid® solution thicknesses (Figure 1h). Besides, full-width-half-maximum (FWHM) analysis depicting the feature width of capillary images at varying depths in different imaging windows was plotted. The FWHMs of the capillary tube without 1% Intralipid® solution were similar in the NIR-II, NIR-IIa, and NIR-IIb window (291±13.7 μm, 258.3±9.1 μm and 240.6±4.8 μm, respectively). Whereas the FWHM in the NIR-II and NIR-IIa windows was 4.7-fold and 1.2-fold larger than that in the NIR-IIb window when the penetration depth increased to 5 mm (Figure 1i). These results indicated that the fluorescence of L1057 NPs in the NIR-IIb window might be the best detection window on account of the reduced light scattering and near-zero autofluorescence in the NIR-IIb window.

### 2.2. *In Vivo* Fluorescence Whole-Body Imaging in the NIR-II and NIR-IIb window

To verify the capability of L1057 NPs as a NIR-IIb contrast agent for *in vivo* fluorescence imaging, we performed whole-body imaging of mice in the NIR-II and NIR-IIb window, respectively. After intravenous injection of L1057 NPs (200 μL, 1 mg mL^-1)^, the nude mice were anesthetized with pentobarbital at various time points and imaged by a NIR-II fluorescence whole-body imaging system. Intense fluorescence signals could be distinguished in blood vessels for as long as 9 h (Figure 2a), illustrating their long circulation time *in vivo* as well as excellent chemical stability. After plotting the cross-sectional intensity profiles of the same capillary, we analyzed the SBR and capillary width (based on the FWHM of a Gaussian function fitted to the cross-sectional intensity profile). As shown in Figure 2b and 2c, the capillary vessel diameter was determined to be 0.321 mm and 0.183 mm in NIR-II and NIR-IIb window, respectively. In addition, the SBRs in NIR-IIb window at different time points were remarkably higher than those in NIR-II window (Figure 2d). The low energy (120 mW cm^-2^) and short exposure time used for NIR-IIb imaging indicated that L1057 NPs are an ideal candidate of NIR-IIb contrast agents attributed to their high brightness..These results also indicated that NIR-IIb might be the most appropriate imaging window for long-term *in vivo* angiography and related tracing, for fluorescence imaging in the NIR-IIb window can achieve higher spatial resolution and SBR than that in the NIR-II window.

**Figure 2.**
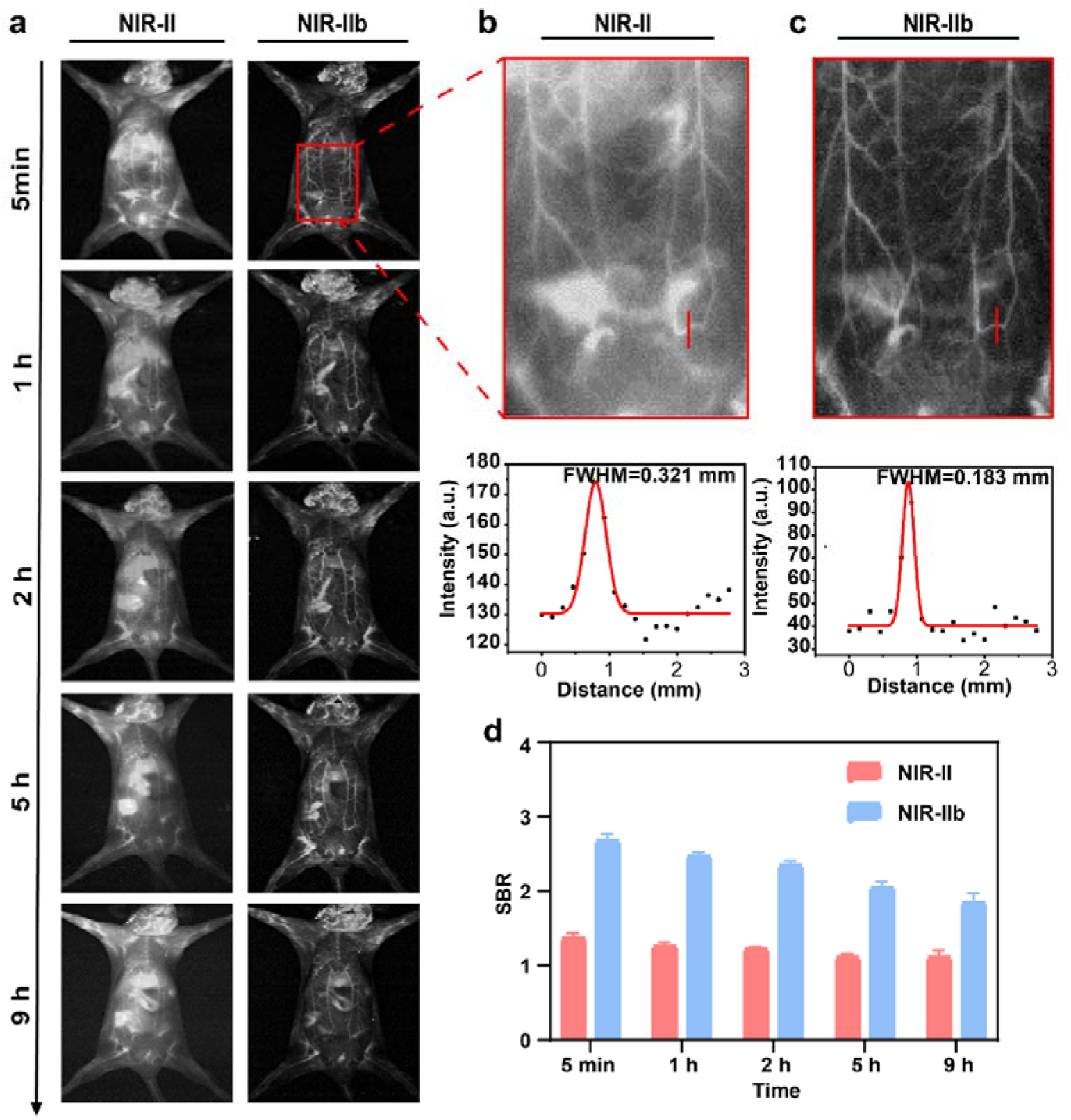
Fluorescence imaging of whole-body blood vessels in living mice with different LP filters using L1057 NPs as contrast agent. (a) The whole-body blood vessel imaging in NIR-II window (1000 nm LP, 20 mW cm^-2^, 10 ms) and NIR-IIb window (1500 nm LP, 120 mW cm^-2^, 150 ms) at different time points. (b) The zoomed-in images of abdomen (top), rectangled in red, in panel a, and the cross-sectional fluorescence intensity profile fitted with Gaussian along red-dashed line (bottom). (c) SBR analysis of the same vessel at different time points in NIR-II and NIR-IIb window.

### 2.3. Real-time Imaging of the Hindlimb Ischemic Reperfusion in the NIR-II and NIR-IIb window

Ischemia is frequently induced in fractures and other peripheral arterial diseases.^[12]^ After ectopic replantation and functional reconstruction, real-time monitoring of ischemic reperfusion is critical to evaluate the degree of recovery. To evaluate the capability of L1057 NPs to track the ischemic reperfusion in limbs, we established a hindlimb ischemia model by ligation of the femoral vein and artery using serrefines for 2 h.^[13]^ Figure 3a shows the experimental schedule. After the i.v. injection of L1057 NPs (200 μL, 1 mg mL^-1^) into the mice, the serrefines on left hindlimb and right hindlimb were sequentially removed and the corresponding ischemic-reperfusion processes were imaged continuously in the NIR-II and NIR-IIb window, respectively. As shown in Figure 3b, the regions of interest were marked with red squares. The ischemic-reperfusion processes could be accurately recorded in the NIR-IIb window but was indistinct in the conditional NIR-II window because of the high background signal interference induced by ischemia-reperfusion injury. Further SBR analysis of the end of hindlimb vessels (EHV) and the reperfusion ratio of hindlimb vessels (HVR) also demonstrated that it was the NIR-IIb window not the conventional NIR-II window could accurately visualize the process of the vessel reperfusion (Figures 3c and 3d), which is crucial for the assessment of blood vessel and tissue damage and subsequent therapy. Taken together, NIR-IIb imaging performed with L1057 NPs could be applied to trace instantaneous blood reperfusion in living mice with real-time feedback and enhanced SBR.

**Figure 3.**
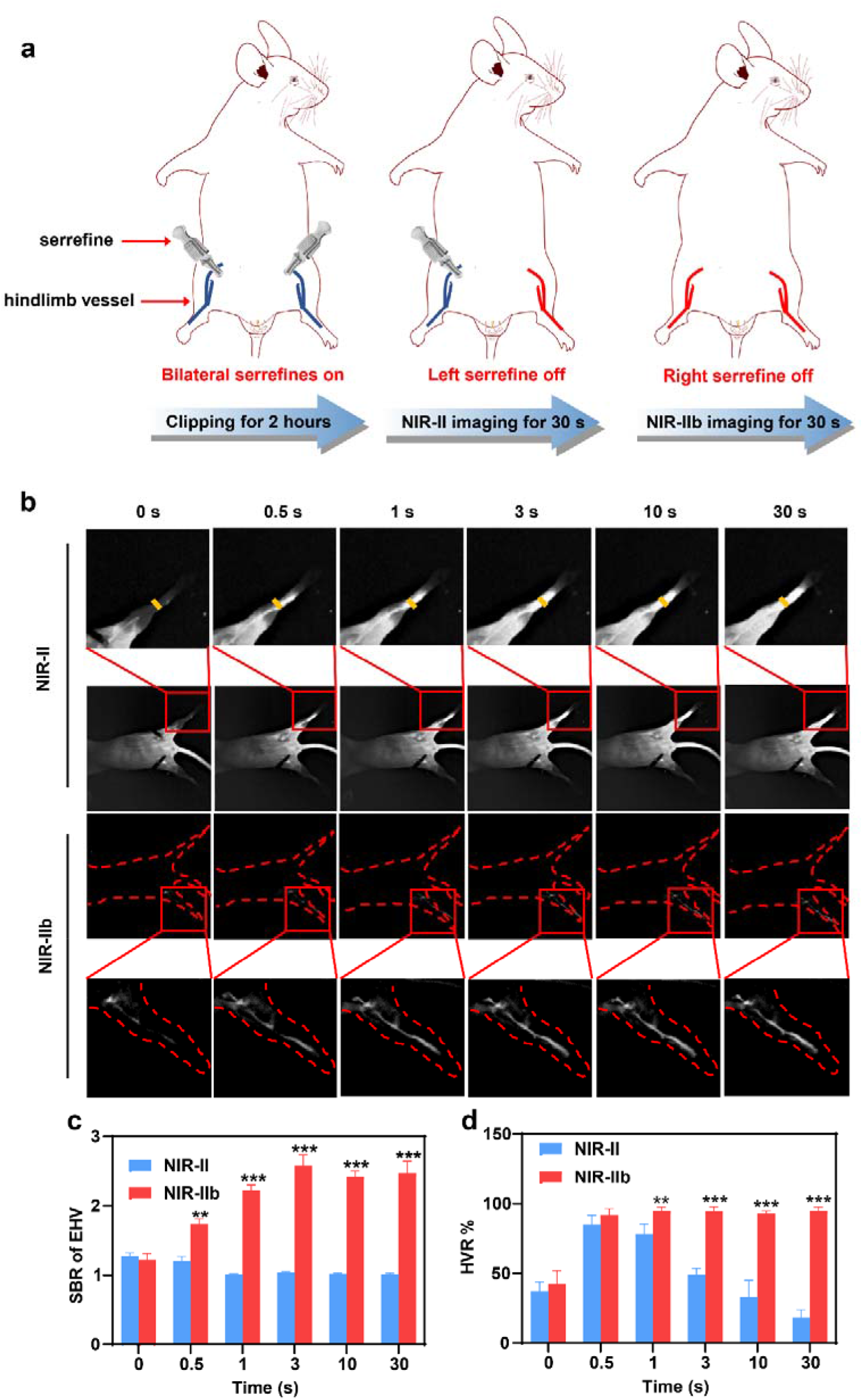
Real-time fluorescence imaging of the ischemic reperfusion in hindlimb. (a) The hindlimb ischemic reperfusion model establishment schematic diagram. (b) NIR-II and NIR-IIb imaging of ischemic reperfusion process at different time points after clipping treatment as indicated. The ischemic reperfusion process was amplified. Yellow arrows indicate the flow front. Scale bar = 10 mm. (c) The SBR analysis of EHV and (d) The HVR analysis at different time points in NIR-II and NIR-IIb windows.

### 2.4. Noninvasive and Invasive Cerebrovascular Imaging in the NIR-II and NIR-IIb window

Emerging evidence indicated that undue vessel growth or abnormal vessel regression actively participate in the pathogenesis of neurological disorders,^[14]^ the deciphering of cerebrovasculature was essential to understand the specific pathogenesis of cerebrovascular dysfunction-related diseases. In this study, we evaluated the feasibility of NIR-IIb fluorescence wide-field cerebrovascular imaging with L1057 NPs. L1057 NPs (200 µL, 1 mg mL^-1^) were firstly intravenously injected into the noninvasive group, which were kept intact skin and skull. It was shown that NIR-IIb images with a large field revealed clearer and more detailed blood vessels compared with NIR-II images. Gaussian fitting analysis of a middle vessel showed that NIR-IIb fluorescence imaging has a smaller FWHM and higher SBR, compared with images obtained in the NIR-II window (Figure 4a). Further, the micro imaging was also conducted to detect cerebral microvessels in the invasive group with a cranial window in the NIR-II window and NIR-IIb window (Figure 4b) at an imaging depth of 200 μm. Likewise, the Figure 4b showed that the same cerebral vessel of interest (red square) detected in the NIR-IIb window exhibited an obvious advantage over that in the NIR-II window in the terms of SBR and spatial resolution (Figure 4b). These results suggested that NIR-IIb fluorescence should be given priority while conducting cerebrovascular imaging using L1057 NPs.

**Figure 4.**
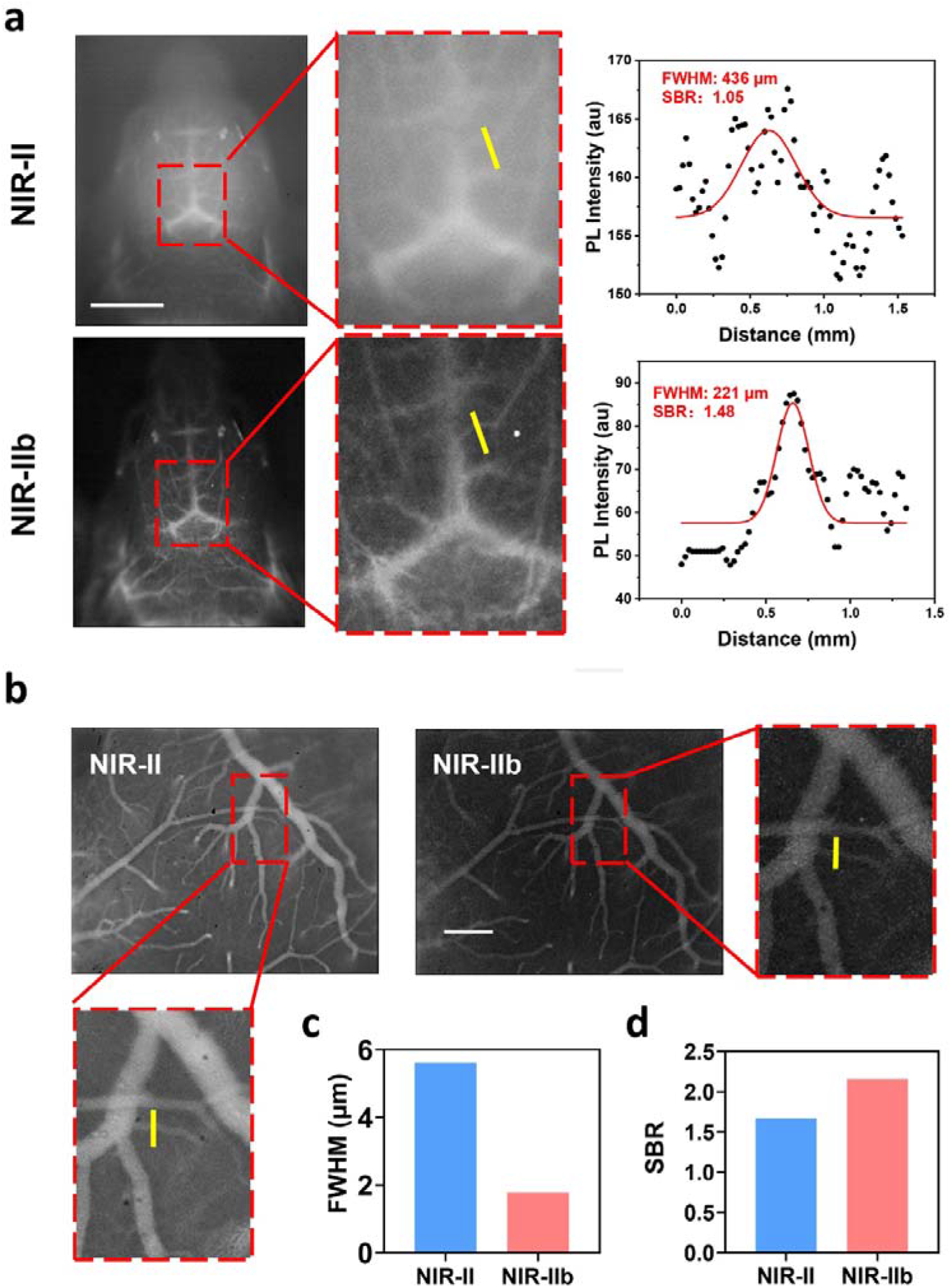
NIR-IIb fluorescence cerebrovascular tomography and functional cerebrovascular imaging. (a) The noninvasive cerebrovascular imaging in the NIR-II and NIR-IIb windows in the macro-imaging system, and the cross-sectional fluorescence intensity profile fitted with Gaussian along red-dashed lines. Scale bar = 5 mm (b) The invasive cerebrovascular imaging in the NIR-II and NIR-IIb using the microscopic imaging system with 5X objective. (c-d) The FWHM and SBR were calculated in the two windows. Scale bar = 1 mm (NA=1.05, Olympus).

### 2.5. Gastrointestinal (GI) Tract Imaging in the NIR-II and NIR-IIb Window

Clinically used GI-imaging techniques like magnetic resonance imaging (MRI) and computed tomography (CT) have unlimited penetration, but their uses for real-time and long-term monitoring of GI are restricted by limited spatial resolution, long acquisition time, and radiation risk to patients and operators.^[7a, 15]^ It is an urgent need for safer, accessible and improved methodology for non-invasive imaging of the GI tract. Fortunately, NIR-IIb fluorescence imaging does not suffer from these drawbacks and is suitable for dynamic intestinal processes visualization. The excellent NIR-IIb imaging capability of L1057 NPs in whole-body and cerebral vascular imaging as well as superior chemical stability in severe acidic conditions (e.g., pH = 1) demonstrated in our previous study encouraged us to evaluate their ability to monitor GI functions.^[10]^ After L1057 NPs (50 µL, 1 mg mL^-1^) were orally injected into healthy nude mice, GI tract imaging was carried out at various time points using different LP filters (1000 and 1500 nm) for fluorescence signal collection. As displayed in Figure 5a, the stomach-duodenum, jejuno-ileum, and colon could be visualized after gavage at 5 min, 5, 9 and 20 h in sequence. Although the GI tract could be detected in the conventional NIR-II window, the images were blurry with low spatial resolution. On the contrary, clear resolution of tissue features was distinguishable with a negligible background in the NIR-IIb window, which were further illustrated in the zoomed-in images of GI tract (Figure 5b). In addition, representative cross-sectional intensity profiles of one of the capillary in Figure 5b showed Gaussian-fitted FWHM of 3.53 mm (SBR = 5.23) in the NIR-IIb window and 5.98 mm (SBR = 1.81) in the NIR-II window (Figure 5c). Thus, L1057 NPs can serve as a NIR-IIb fluorescence contrast agent to visualize deep tissues and could be an excellent platform for non-invasively assessing GI tract-related diseases.

**Figure 5.**
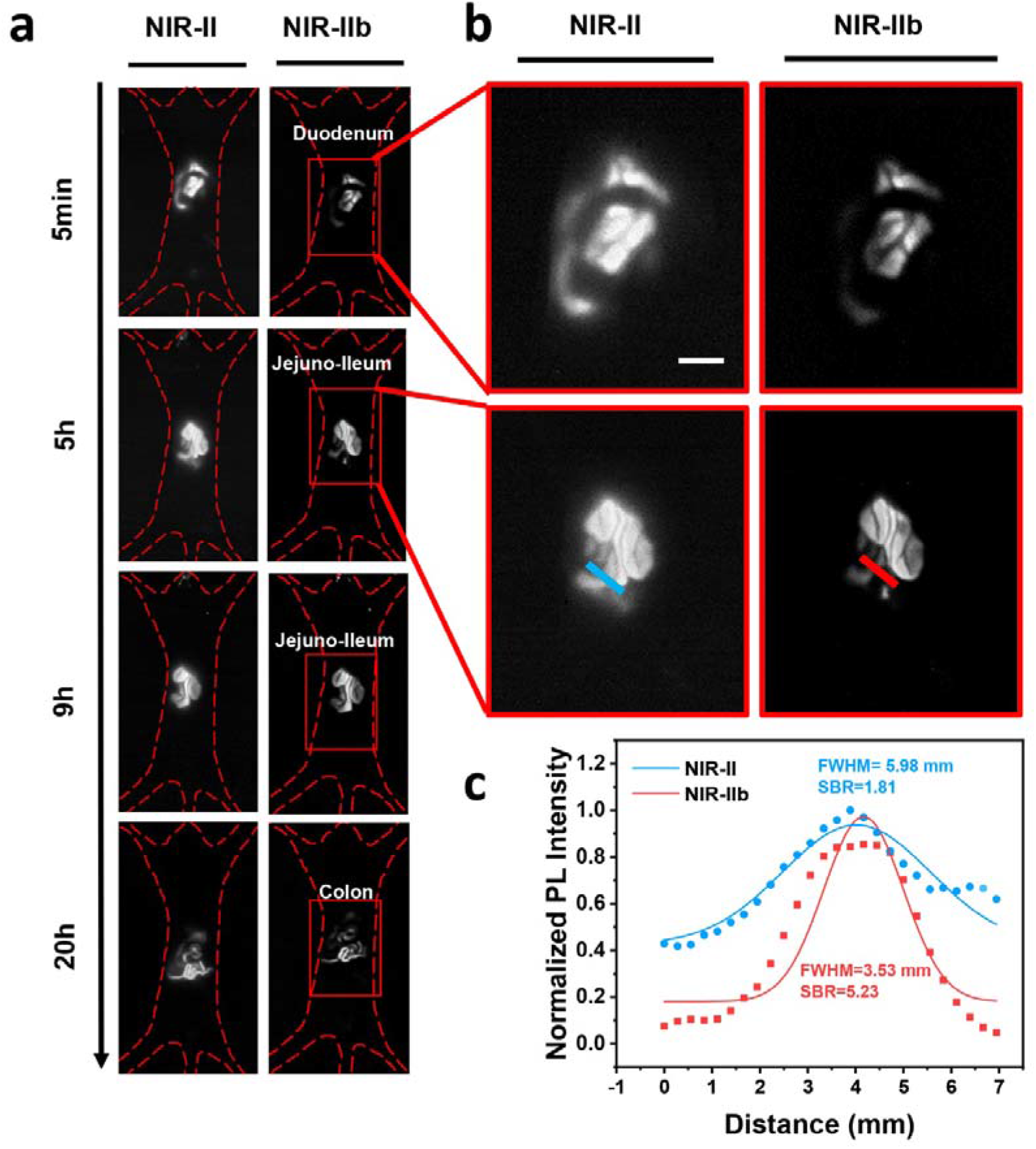
Fluorescence imaging of the GI tract. (a) Real-time monitoring of intestinal peristalsis in living mice gavaged with L1057 NPs (300 μL, 1 mg mL^-1^) using different LP filters (1000 nm LP, 10 ms, 20 mW cm^-2^; 1500 nm LP, 150 ms, 100 mW cm^-2^) at various time points (5 min, 5, 9 and 20 h). (b) Representative images in the NIR-II and NIR-IIb window at the same time point, and (c) the FWHMs and SBRs were analyzed in the two windows.

### 2.6. Tumor Progression Bioimaging in the NIR-II and NIR-IIb Window

Early diagnosis and tracing of tumor *in vivo* are of great importance since they can help us well understand the progression of tumors.^[16]^ Previous study has demonstrated that L1057 NPs exhibited remarkable tumor marking capability attributed to the enhanced permeability and retention (EPR) effect.^[10]^ To further assess the ability of L1057 NPs for tumor progression monitoring in different stages, we established xenograft renal cell carcinoma (RCC) models by subcutaneous injection of OSRC2 cells into the foreleg armpit of the male BALB/c nude mice, and controlled the time of tumor growth to mimic the early-, mid- and late-stage of tumor (Stage I: 2 weeks of tumor formation, that is, the initial tumor formation; Stage II: 4 weeks of tumor formation, significantly enlarged tumor with a volume> 200 mm^3^; Stage III: tumor formation time > 6 weeks, the tumor continued to grow, and the mouse appears cachexia (skin and bone, extremely thin, and other typical late symptoms)). Following the intravenous injection of L1057 NPs (200 μL, 1 mg mL^-1^) through the tail vein, fluorescence imaging of tumors was performed 2 h post injection. As shown in Figures 6a-c, bright fluorescence signals can be clearly detected from the tumor sites due to the EPR effect. The vascular network patterns became gradually complicated with increased vascular density from stage I to III. Higher spatial resolution was obtained in the NIR-IIb window than in the NIR-II window, benefited from the advantages of NIR-IIb fluorescence imaging. Similarly, the tumor to normal tissue signal ratio (T/TN) was also significantly higher in the NIR-IIb window than that in the NIR-II window (Figures 6d-f). It was noted that T/NT value became increased from stage I to III regardless the imaging window, indicating that L1057 NPs are more likely to accumulate in advanced tumors.

**Figure 6.**
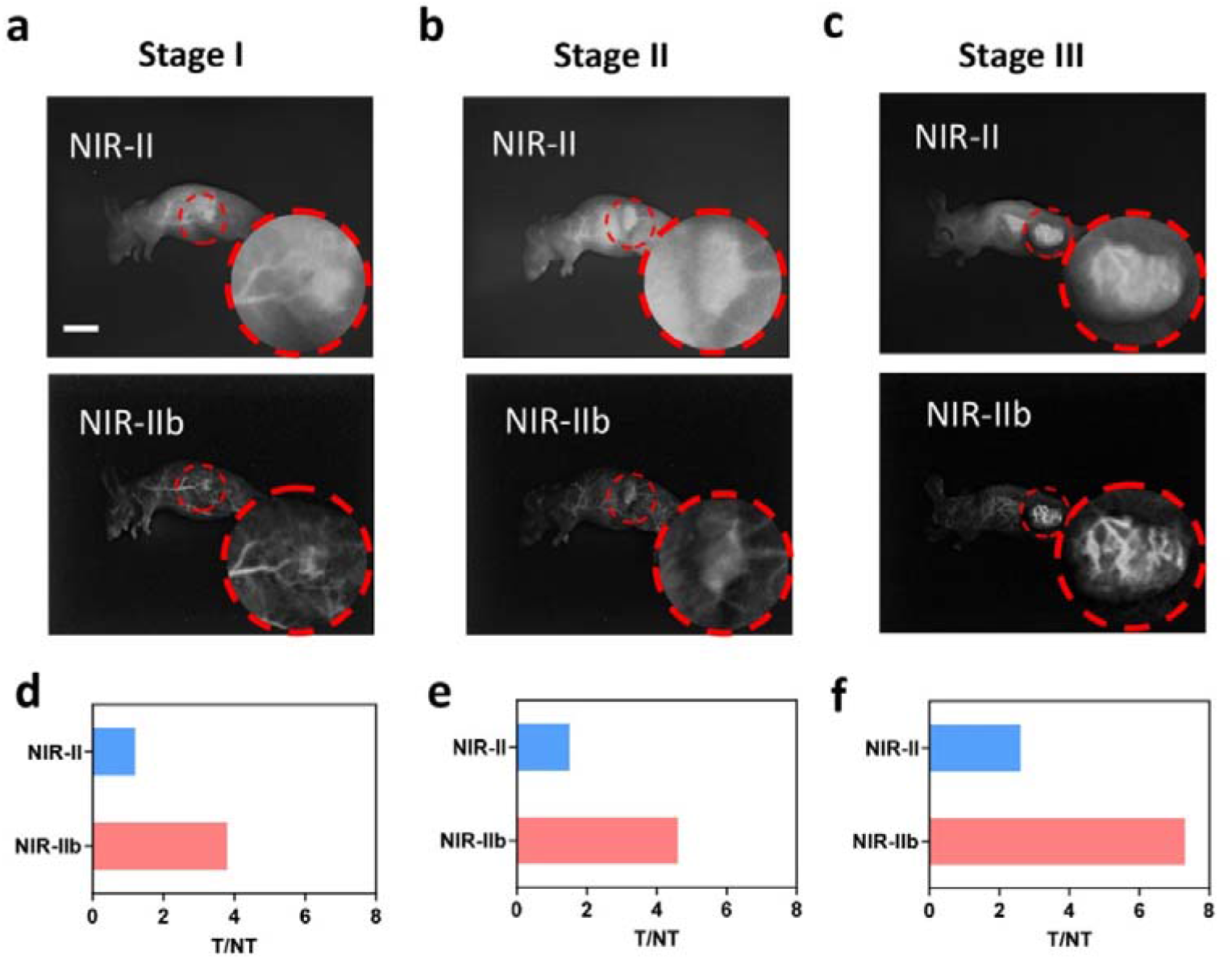
Fluorescence imaging of tumor progression in different NIR-II subwindows. (a-c) Representive images of subcutaneous tumor-bearing mice at different tumor stages after intravenously injected with L1057 (200 μL, 1 mg mL^-1^) in NIR-II and NIR-IIb windonw. Laser power: 20 mW cm^-2^ and 120 mW cm^-2^ for NIR-II and NIR-IIb imaging, respectively. Exposure time: 10 ms and 150 ms for NIR-II and NIR-IIb imaging, respectively. Scale bar: 10 mm. (d-f) Tumor-to-normal tissue ratio based on the fluorescence signal.

It is well-known that the formation and growth of a tumor are highly dependent on the blood supply around the tumor. Therefore, noninvasive imaging of tumor vessels with high spatial resolution may provide more detailed information on tumor vessel morphology, length, and leakage characteristics.^[17]^ Importantly, vascular imaging in a real-time manner that equips both anatomic and hemodynamic information will extremely facilitate cancer detection and accurate assessment of therapeutic effects.^[18]^ In this study, the tumor vessels displayed more details in the NIR-IIb window than those in the NIR-II window, which may contributed to better understand the cancer status. Together, the NIR-IIb window might be the most appropriate imaging window for L1057 NPs in tumor imaging to better understand the progression of tumors.

## 3. Conclusion

In summary, for the first time, we demonstrated the potential of a highly bright SPN of L1057 NPs as an *in vivo* NIR-IIb fluorescence contrast agent, with excellent biocompatibility, high NIR-IIb brightness, and high stability. *In vitro* and *in vivo* screening experiments revealed that excitation at longer wavelength and avoiding the overtone peak of water is beneficial to achieve higher SBRs and deeper tissue penetration. Specifically, 915 nm is an appropriate excitation wavelength for L1057 NPs to realize high quality imaging. Benefiting from significantly suppressed photon scattering and nearly zero autofluorescence in the NIR-IIb window, we further demonstrated in various animal models that L1057 NPs as fluorescence contrast agent afforded better spatial resolution, SBR and penetration depth than those in the NIR-II window. This work not only illustrates that L1057 NPs are a promising candidate material for NIR-IIb contrast agent, but also suggests that simultaneous optimization of excitation wavelength and emission is an efficient strategy to improve imaging quality. Future work will focus on the development of novel high brightness and/or activated NIR-IIb SPNs with longer excitation wavelength for *in vivo* fluorescence imaging.

## 4. Experimental Section

### Materials

1,2-Disteroyl-*sn*-glycero-3-phosphoethanolamine-*N*-[methoxy(polyethylene glycol)-2000](DSPE-PEG_2000_) was purchased from Laysan Bio Inc. Milli-Q water (18.2 MΩ) was obtained through a Milli-Q Plus system (Millipore Corporation, Bedford, USA) and was utilized for all the experiments involving aqueous medium. The fluorophore indocyanine green (ICG) was purchased from DanDong Pharmaceutical Factory (Liaoning, China). All other chemicals were obtained from Sigma-Aldrich or Energy Chemical (China) and were used as received without further purification.

### Absorption and Photoluminescence Spectra Characterization

Absorption spectra were recorded on a Shimadzu UV-1750 spectrometer. Photoluminescence (PL) spectra were recorded on an Edinburgh instrument FLS980 equipped with a Xe lamp as an excitation source and a nitrogen-cooled InGaAs detector.

### Intralipid® Phantom Assay

*In vitro* testing in an Intralipid® phantom was performed as described previously. 1% Intralipid® solution was prepared by diluting 20% Intralipid® into Milli-Q water. A capillary glass tube (Inner diameter = 0.3 mm) filled with L1057 NPs aqueous dispersion (1 mg mL^-1^) was immersed in the prepared 1% Intralipid® solution, the depth of which ranged from 1 to 6 mm below the top surface. NIR-II and NIR-IIb imaging at different depths were performed.

### Optical setup for NIR-II Fluorescence Macro-Imaging System

A 2D electronic-cooling InGaAs camera (640 pixels × 512 pixels, TEKWIN SYSTEM, China) equipped with a prime lens (focal length: 35 mm, antireflection coating at 900-1700 nm, TEKWIN SYSTEM, China) was utilized to collect and acquire the fluorescence signals with different filters. A 915 nm laser beam was coupled to a collimator and expanded by a lens to provide uniform illumination on the field. The facular power density was measured and adjusted before performing each experiment (see Figure S3). All the LP filters were purchased from Thorlabs (USA).

### Optical Setup for NIR-II Fluorescence Mesoscopic/Microscopic Imaging System

*The NIR-IIb fluorescence microscopic imaging optical system (NIRII-MS, Sunnyoptical) was equipped with a 1080 nm long-pass dichroic mirror (DMLP) and objectives. The tube lens had 1000–1700 nm anti-reflection. A 915 nm laser was utilized as an excitation source. After 915 nm laser beams reflect from a 1080 nm DMLP and pass through an objective, the observed sample is illuminated. The objective could be an air objective lens (LSM03, WD = 25*.*1 mm, Thorlabs or LMS05, WD = 36 mm, Thorlabs), or an infrared transmission water-immersed object (XLPLN25XWMP2, 25 ×, NA = 1*.*05, Olympus). NIR-II fluorescence mesoscopic/microscopic images of the sample were collected with the InGaAs camera after passing through different long-pass filters (Thorlabs) (see Figures S5 and S6)*.

### Animal Experiments

Institute of Cancer Research (ICR) mice (6–8 weeks old, female) and BLAB/c Nude mice (6–8 weeks old, female) were provided from the SLAC laboratory Animal Corporation (Shanghai, China) and kept in the Laboratory Animal Center of Zhejiang University (Hangzhou, China). The animal housing area was maintained at 24 °C with a 12 hours light/dark cycle, with free water and food available. Before each operation and imaging experiment, mice were anesthetized via intraperitoneal injection of 2% pentobarbital (40–50 mg kg^-1^) and kept in maintaining anesthesia. Mice were intravenously injected with L1057 NPs (0.01 mg g^-1^ body weight, i.v.) before imaging.

### In Vivo Fluorescence Whole-Body Imaging

Nude mice (female, ∼20g) were chosen for NIR-II fluorescence whole-body imaging. After intravenous injection of L1057 NPs (200 μL, 1 mg mL^-1^), the mice were anesthetized with pentobarbital continuously and stably, and imaged with the NIR-II fluorescence macro-imaging system. The conventional NIR-II images were collected with excitation at 915 nm (20 mW cm^-2^) and signal collection using a 1000 nm LP filter. The exposure time was 10 ms. The NIR-IIb imaging was achieved under 915 nm laser excitation (120 mW cm^-2^) using a 1500 nm LP filter and the exposure time was 150 ms.

### Fluorescence Imaging of Ischemic Reperfusion in Hindlimbs

Nude mice (female, ∼20g) were anesthetized with pentobarbital. In the anesthetized state, bilateral hindlimb ischemia was induced by ligation of the left vein and femoral artery using vascular clamps for 2 h. The conventional NIR-II imaging was performed on the left ischemic hindlimbs while the NIR-IIb imaging on the right hindlimbs. For dynamic imaging, the camera was set to operate continuously. Subsequently, L1057 NPs (1 mg mL^-1^, 200 μL) were injected into the mice through the tail vein and the clips were removed and the ischemic reperfusion in hindlimbs was evaluated for 30 s.

### In Vivo Fluorescence Cerebrovascular Imaging

*In vivo* fluorescence cerebrovascular imaging was performed using fluorescence macro/microscopic imaging optical system. The mice were divided into two groups (noninvasive and invasive). The noninvasive group was kept intact skull and skin^[1a]^ while the invasive group was performed microsurgery to remove the cranial window.^[11b]^ The noninvasive NIR-II and NIR-IIb fluorescence cerebrovascular images were recorded using the NIR-II fluorescence micro-imaging system immediately after L1057 (1 mg mL^-1^, 200 µL) was injected through tail vein (Imaging conditions for noninvasive imaging: 915 nm laser power density = 20 mW cm^-2^ (NIR-II), and 120 mW cm^-2^ (NIR-IIb); exposure time = 10 ms (NIR-II), and 150 ms (NIR-IIb)). The invasive NIR-II and NIR-IIb fluorescence cerebrovascular images were obtained using the NIR-II fluorescence microscopic imaging system (Imaging conditions for noninvasive imaging with 5X objective: 915 nm laser power density= 60 mW cm^-2^ (NIR-II), 250 mW cm^-2^ (NIR-IIb); exposure time = 10 ms (NIR-II), and 150 ms (NIR-IIb)).

### Tumor Model Establishment

The OSRC2 cells (human renal cell carcinoma cell) were obtained from the Cell Culture Center of the Institute of Basic Medical Sciences, Chinese Academy of Medical Sciences (Shanghai, China) and cultured in the standard media recommended by American type culture collection (ATCC), and corresponding tumor models were established via the subcutaneous injection of OSRC2 cells (1×10^10^ cells in 100 μL serum-free medium) into the foreleg armpit of the male BALB/c nude mice.

### Data Analysis

Quantitative analysis of each fluorescent image was performed based on the measurement of mean signal intensity in the manually selected regions of interest, using Image J software (Version 1.6.0, National Institutes of Health, USA). Graphs were generated using Origin Pro software (Version 9.0; OriginLab Company, US).

## Supporting Information

Supporting Information is available from the Wiley Online Library or from the author.

## Acknowledgements

D. Xue, H. Zhou and Z. Lu contributed equally in this work. This work was supported by the National Natural Science Foundation of China (81672508, 81672520, 81870484, 61975172 and 82001874), Science and Technology Planning Project of Zhejiang (2019C03089), Zhejiang Medical and Health Science and Technology Plan Project (WKJ-ZJ-2031), Zhejiang Provincial Natural Science Foundation of China (LR17F050001), the NanjingTech startup Grant (38274017115), Jiangsu Provincial Foundation for Distinguished Young Scholars (BK20170041), Open Research Fund of Anhui Key Laboratory of Tobacco Chemistry (20181140), China-Sweden Joint Mobility Project (51811530018), Natural Science Foundation of Zhejiang Province (LY21F050005), and The Fundamental Research Funds for the Central Universities (2021FZZX005-18).

## Conflict of Interest

The authors declare no conflict of interest.

## Notes

### Competing Interest Statement

The authors have declared no competing interest.

