## Supporting Information for "Simultaneous Optimization of Excitation Wavelength and Emission Window for Semiconducting Polymer Nanoparticles to Improve Imaging Quality"

Dr. D. Xue, Z. Lu, Dr. A. Zebiluba, Prof. J. Qian, Prof. G. Li

Department of Urology, Sir Run-Run Shaw Hospital, School of Medicine, Zhejiang University, Hangzhou, 310016, China.

H. Zhou, M. Li, L.Li, Prof. J. Liu

Key Laboratory of Flexible Electronics (KLOFE) Institute of Advanced Materials (IAM), Nanjing Tech University, Nanjing 211800, China.

Y. Zhang, Z. Feng, Prof. J. Qian

State Key Laboratory of Modern Optical Instrumentations, Centre for Optical and Electromagnetic Research, College of Optical Science and Engineering, International Research Center for Advanced Photonics, Zhejiang University, Hangzhou, 310058, China.

Dr. Z. Xu

Institute of Intelligent Optoelectronic Technology, Zhejiang University of Technology, Hangzhou, 310014, China

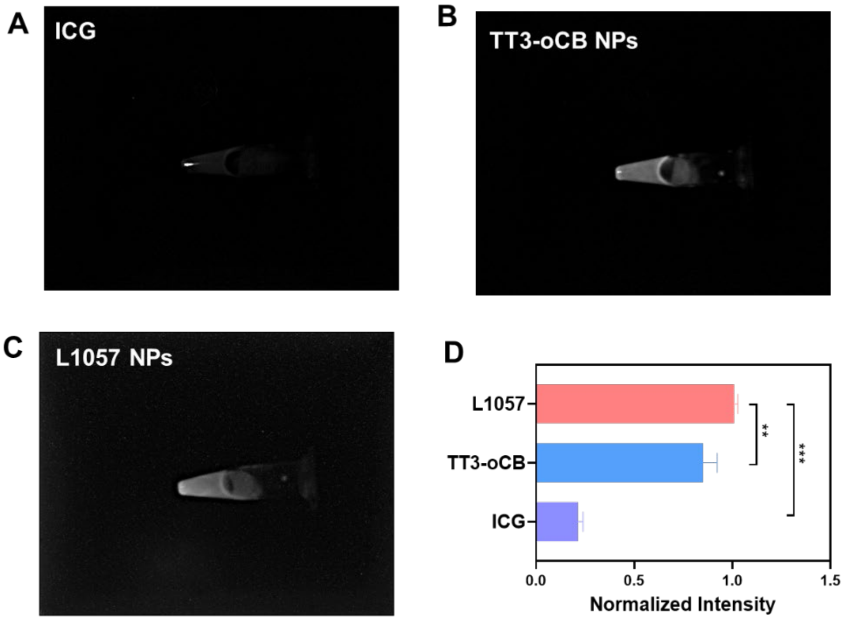


**Figure S1.** The fluorescence intensity comparison of ICG, TT3-oCB NPs and L1057 NPs at the same concentration (0.05 mg $mL^{-1}$)in the NIR-IIb window (ICG and TT3-oCB NPs were excited by 793 nm laser, L1057 NPs were exited by 915 nm laser; Power Intensity: 20 mW $cm^{-2}$, 100 ms).

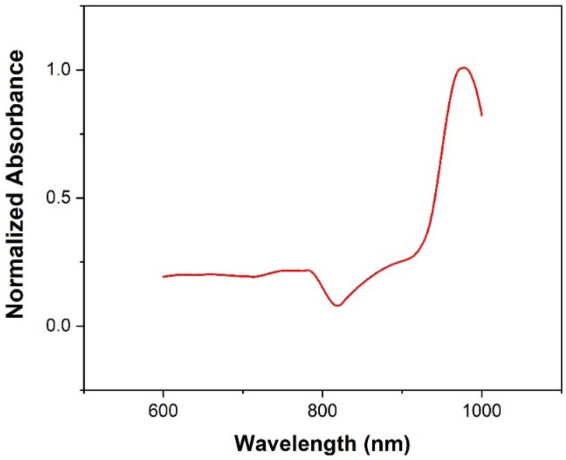


**Figure S2.** The water absorption spectrum (600-1000 nm)

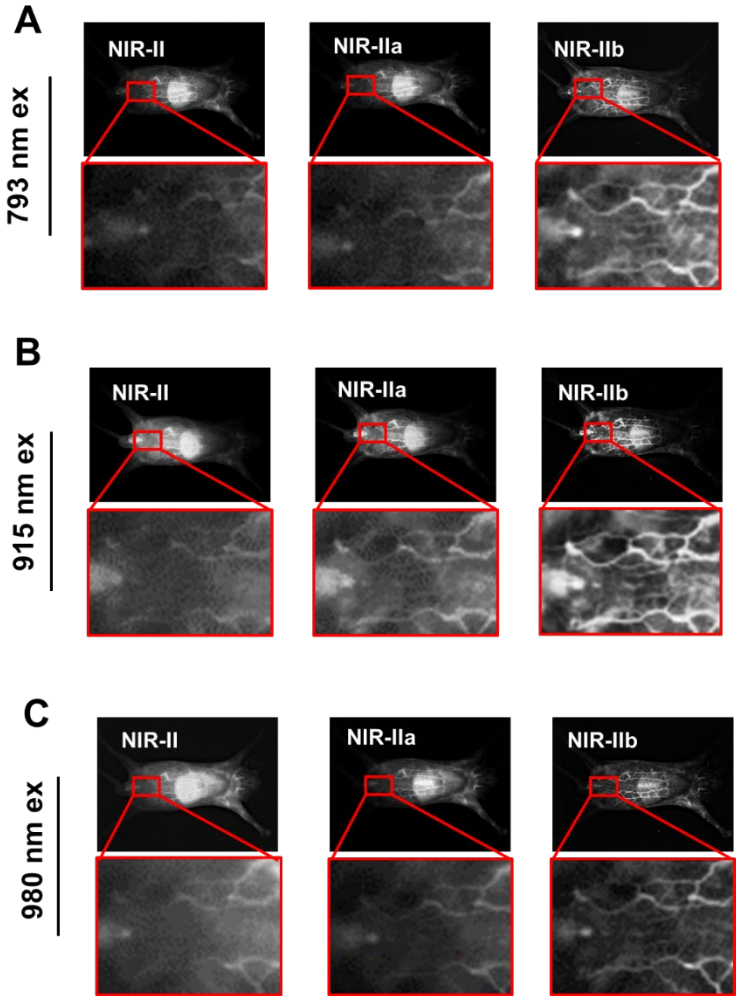


**Figure S3.** The whole-body imaging of L1057 NPs in NIR-II, NIR-IIa and NIR-IIb windows excited by 793 nm laser, 915 nm laser and 980 nm laser, respectively.

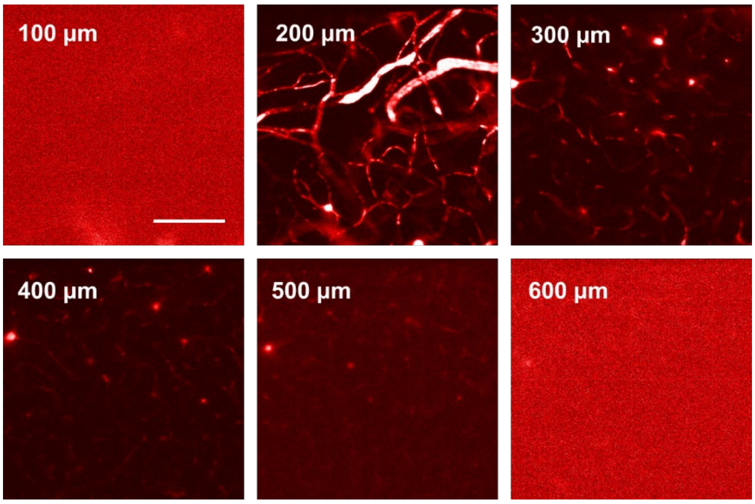


**Figure S4.** NIR-II fluorescence confocal microscopic in vivo imaging of cerebral blood vessels of the mouse at 6 typical depths (100 μm, 200 μm, 300 μm, 400 μm,500 μm and 600 μm).

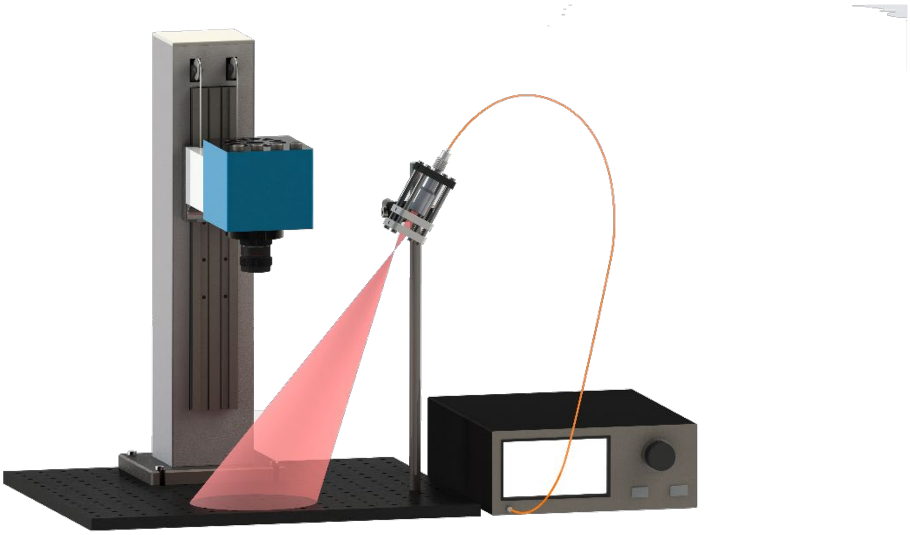

**Figure S5.** Schematic illustration for NIR-II fluorescence macro-imaging system

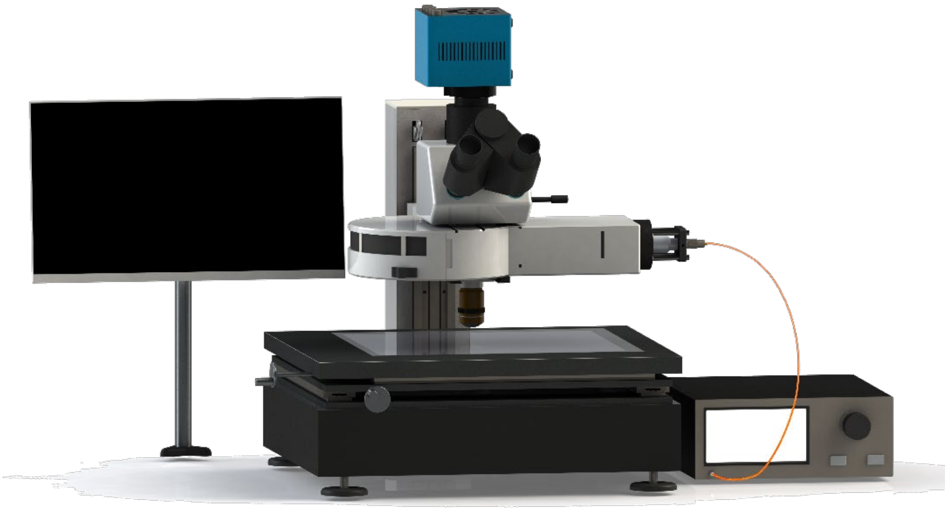

**Figure S6.** Schematic illustration for NIR-II fluorescence micro-imaging system
